# Maternal separation on the choroid plexus of UChA and UChB rats

**DOI:** 10.1101/2025.11.27.690964

**Authors:** M. Martinez, N. G. S. Sidani, F. E. Martinez

**Affiliations:** Department of Morphology and Pathology, Federal University of São Carlos (UFSCar), São Carlos, SP, Brazil; Department of Structural and Functional Biology, State University of São Paulo (UNESP), Botucatu, SP, Brazil

**Keywords:** Ethanol, Neonatal stress, Plexus chorioid, UChA and UChB rats

## Abstract

Alcoholism is one of the oldest and most widespread drug addictions described in the literature and one of the biggest concerns in the health area. There is a growing number of investigations into the medical, social, and economic problems triggered by the abusive consumption of ethanol-based distillates or fermented products that can affect tissues or organs in different ways. Intrinsic or extrinsic situations that threaten organic homeostasis trigger a series of adaptive responses that constitute the stress systems, which consist of different physical and mental reactions that oppose the stressful stimulus, aiming to restore the lost balance. The UChA and UChB rat varieties constitute rare models for studies related to the genetic, biochemical, physiological, nutritional and pharmacological factors of the effects of alcohol, as well as appetite and tolerance, important factors related to human alcoholism. The present work arose from the interest in investigating the effects of stress on these animals, since several aspects of alcoholism may be the result of stress experienced early in life, being potentiated or not in the adult individual. Thus, the research proposes to investigate and evaluate whether maternal neonatal separation, applied to male offspring of UChA and UChB rats, potentiates the toxic effects of chronic alcohol intake on choroid plexus cells. Maternal separation (MS) change the ethanol consumption and not altered IGF-I expressions. There was a negative association between ethanol consumption and MS in body mass gain and other correlations with feed, water consumptions and hormones dosages. Maternal separation potentiated epithelial atrophy of the choroid plexus in UChA and UChB animals.

## INTRODUCTION

Alcoholism is one of the biggest public health problems worldwide. It can be defined as a multifactorial syndrome, with physical, mental, and social impairment (Edwards & Gross, 1976). Ethanol use is common in most cultures, but in varying doses and extents (Wolfe *et al*. 2023). There are few studies with a morphological focus on the effects of ethanol consumption on the choroid plexuses. Changes in the mouse fetal choroid plexuses following chronic maternal alcoholization were observed by Craciun *et al*. (1989). Tirapelli *et al*. (2000) described ultrastructural lesions in the choroid plexuses of the lateral ventricles of rats subjected to ingestion of 30 GL sugar cane spirit for 300 days. Immunofluorescence studies reported reduced glucose conductors in the choroid plexus of ethanol-treated Sprague-Dawley rats (Handa *et al*. 2000).

Fetal alcohol spectrum disorder encompasses the range of deleterious outcomes of prenatal alcohol exposure in the affected offspring, including developmental delay, intellectual disability, attention deficits and conduct disorders (Sambo & Goldman, 2023). Fetal alcohol exposure leads to severe disruptions in learning and memory involving the hippocampus and prefrontal cortex in humans (Heroux *et al*. 2018)

Intrinsic or extrinsic situations that threaten organic homeostasis trigger a series of adaptive responses that constitute stress systems, which consist of different physical and mental reactions, which oppose the stressful stimulus, aiming to reestablish the lost balance. Responses to stress are characterized by behavioral and physical changes, coordinated by the integration of the central nervous system with peripheral systems (Levine, 2000).

Early life experiences are one of the factors affecting psychological and physiological development and may lead to significant alterations in emotion and stress responses in later life (Protsky *et al*. 2005). Childhood adversity can induce maladaptive behaviors and increase risk for affective disorders, post-traumatic stress disorder, personality disorders, and vulnerability to stress in adulthood. Early environmental stress significantly affects the development of offspring and has been modeled in rats through the maternal separation paradigm, which alters the functioning of the HPA axis and can enhance ethanol intake at adulthood (Pautassi *et al*. 2012). Deprivation of maternal care interrupts brain development through the disturbance of various neurotransmitters; however, the details remain unclear (Mavrenkova *et al*. 2023). Neonatal maternal separation (MS) has been used to model long-lasting changes in behavior caused by neuroplastic changes associated with exposure to early-life stress (Hasegawa *et al*. 2023).

Rats subjected to neonatal manipulation, when subsequently exposed to aversive stimuli, show a smaller increase in plasma ACTH and corticosterone than control rats, in addition to a faster return of these hormones to basal concentrations. The origin of this response appears to be permanent changes in central regulatory mechanisms – neonatal manipulation stimulates the release of thyroid-stimulating hormone which appears to stimulate serotonergic projections within the hippocampus. Serotonin can cause a long-term increase in the number of glucocorticoid receptors in the hippocampus, resulting in increased sensitivity of the region to circulating glucocorticoids and, consequently, increasing the effectiveness of the negative feedback mechanism, which reduces post-stress secretion of ACTH and corticosterone (Levine, 2000).

Insulin homologous growth factor (IGF) is a polypeptide responsible for functional homeostasis in different tissues. It is reported structurally as a mitogenic factor, with multiple neurotrophic actions in the nervous system and in the adult central nervous system. IGF is a neuromodulator of the adult brain, acting as a pre- and post-synaptic messenger. Glial cells are likely involved in the actions of IGF (Garcia Segura *et al*. 1997). Levels of growth factors homologous to insulin vary with age and may be selective in brain areas. Fetal ethanol ingestion during pregnancy is the leading cause of preventable cognitive impairment. Maternal EtOH exposure increased mitochondrial DNA damage (mtDNA) in fetal brain tissue and IGF-1 rescued neurons from EtOH-mediated mtDNA damage and OGG1 inhibition (Darbinian *et al*. 2023).

The UChA and UChB rat varieties constitute rare models for studies related to the genetic, biochemical, physiological, nutritional and pharmacological factors of the effects of alcohol, as well as appetite and tolerance, important factors related to human alcoholism. These models of experimental alcoholism were selected from *Rattus norvegiccus albinus* during the 1950’s at the University of Chile, from which the name UCh originated. UChA rats have a low voluntary consumption profile (0.1-2 grams of ethanol per kilogram of body weight per day) and UChB rats have a high voluntary consumption profile (5-7 grams of ethanol per kilogram of body weight per day) (Mardones & Segovia Riquelme, 1983). The lineages are close to the chronic human dependence on ethanol reality since the animal’s present voluntary ethanol consumption. Thus, interest was aroused in investigating the effects of stress on these animals because several aspects of alcoholism may be the result of stress experienced early, being enhanced or not in the adult individual. The research proposes to analyze and evaluate whether neonatal maternal separation, applied to male offspring of UChA and UChB rats may potentiates the toxic effects of chronic alcohol intake on choroid plexus cells.

## MATERIAL AND METHODS

### Animals

The UChA strain rats (low voluntary 10 % ethanol consumption), UChB (high voluntary 10% ethanol consumption) and Wistar strain were obtained from the Anatomy Department Animal Facility at the Institute of Bioscience, Campus of Botucatu (IBB/UNESP). Four couples of each strain were used with two experimental groups (Maternal separation absence - MSA and Maternal separation - MS). Adult females, after fertilization, were observed during gestational period. The birth date was stipulated as day zero, with the litter being standardized with eight pups, by selecting the largest number of males as possible. Each group used fourteen animals for 120 days of experimentation. The rats were kept in polypropylene cages measuring 40X30X15 cm with laboratory-grade pine shavings as bedding, under controlled conditions of brightness (12 h light and dark cycle), temperature (20 to 25 °C) with water and standard pellet food (Nuvital^®^) *ad libitum*. All animal experiments were performed under a protocol approved by the local ethical committee (IBB/UNESP) and in accordance with the Brazilian Animal Protection Legislation. Six groups were formed:

| UChA | UChA MS | UChB | UChB MS | Control | Control MS |
| --- | --- | --- | --- | --- | --- |
| Ethanol consumption lower than 2g of ethanol/kg body weight/day | Ethanol consumption lower than 2g of ethanol/kg body weight/day | Ethanol consumption higher than 2g of ethanol/kg body weight/day | Ethanol consumption higher than 2g of ethanol/kg body weight/day | Free consumption of water and ration | Free consumption of water and ration |
| Maternal separation absence (MSA) | Maternal separation (MS) | Maternal separation absence (MSA) | Maternal separation (MS) | Maternal separation absence (MSA) | Maternal separation (MS) |

### Hyperresponsivity induction by Maternal Separation

Pups were separated daily from their mothers during the Stress Hyporesponsive Period (SHRP), between the age of 4 and 14 days old, always at the same time. In MS, pups were isolated for 4 hours in a cage containing eight subdivisions, one for each pup. The animals were kept in a adjacent room at 30°C and over 50% humidity. Afterwards, the pups were brought back to their mothers which had been removed from their respective cages before meeting their litter again. Control pups were not separated from their mothers and they were only subjected to the standard procedures used for laboratory animals. Rats were handled by only one researcher during the whole experiment. Animal facility room was isolated as much as possible from external noise. Researcher’s hands were washed in running water, dried and rubbed on the wood of the litter cage roof to eliminate smells that could make mothers reject their offspring. Male pups were housed in boxes with a minimum of two and a maximum of four animals after weaning to 21 days, avoiding stress by social isolation. Male pups were individualized and received standard procedures for the species at 50 days of age.

### Selection of ethanol consuming animals

At 65 days of age, UChA and UChB rats started receiving a 10% ethanol solution and water *ad libitum* for 15 days, alternating bottle (water and ethanol) positions in the cage during the measurements. Animals presenting the highest (UChB) and the lowest (UChA) ethanol consumption were selected. Ethanol (10%) and water were provided *ad libitum* to UChA and UChB rats until the end of the experiment (120 days). UChA and UChB strains were selected and standardized according to Mardones & Segovia-Riquelme (1983). The same number of animals was selected in ethanol consumption absence rats at random.

### Biological materials

The collection of biological materials was held at 120 days of age according to two distinct procedures: (a) seven animals of each group were subjected to euthanasia by decapitation for evaluation of blood plasma hormonal concentrations. Later the rats underwent craniotomy and brain dissected; b) seven of each group animals were anesthetized with an intraperitoneal injection of pentobarbital (20 mg / kg) and transcardiac infused with 0.9 % saline solution, followed by paraformaldehyde 4 % in sodium phosphate buffer (PBS), 0.1M and pH 7, 4, at 40 °C. Brain was collected to histological routine for morphometric and stereological analyses, weighed and fixed by immersion in Bouin’s solution during 24 h. After washing in the increased ethanol series, the tissue fragments were dehydrated and embedded in paraplastic (Oxford Labware, St. Louis, USA). Blocks were sectioned at 8μm thickness in the LEICA 2145 microtome and stained with Hematoxylin-eosin (HE). The slides were analyzed and captured by digital photomicroscope Axio phot II (Zeiss).

### Morphometric and stereological analysis

The brains fixed in paraformaldehyde 4% were washed in 70% alcohol, dehydrated, diaphanized, included in paraplastic and cut with five micrometers thick. Collections of cross-sectional slides were obtained. The sections were stained with Hematoxylin-Eosin, Masson’s Trichrome and IGFR-1. Histological slides stained with HE was used in morphometric analyses. The height of the choroid plexus epithelium and the area and perimeter of the nuclei of these cells were measured in 15 different slides for each animal on three cuts per slide with five measures per cutting. The slides were analyzed and photographed using a Zeiss Axiostar plus photomicroscope at the Anatomy Laboratory of DMP/UFSCar. The Axio vision Computerized Image Analysis System (Zeiss) was used to measure the variables described.

### Hormonal assays

After decapitation, trunk blood samples were obtained from animals to verify LH, FSH *(Harbor-UCLA,USA)*, Testosterone, DHT and Corticosterone (*Maia, BioChem Immuno Systems, Italia SPA*) hormone levels, and plasma concentration was achieved by double antibody radioimmunoassay method (RIA). All samples were tested in the same assay for each hormone to avoid intraassay variations.

### Immunohistochemistry for IGFR-1

For IGFR-1 expression rabbit antibodies were used (Chemicon) according to manufactures details in 5 animals from each group. Five slides of each brain with 5 slices per slide were stained.

### Statistical analysis

Statistical analysis of the variables described was performed through ANOVA followed by Tukey’s multiple comparison tests to compare group means. Results were expressed as mean ± standard error or deviation of the mean. All tests were performed in two-tail modality and *p*<0.05 values were considered statistically significant.

## RESULTS

### Choroid plexus histology

The choroid plexus located in the cerebral ventricles is formed by tufts of capillaries lined by specialized secretory epithelium and connective tissue. Epithelial cells are predominantly cuboidal, with eosinophilic cytoplasm, and have an ovoid nucleus containing delicate and regular chromatin. This simple cubic epithelium forms numerous villi within the cerebral ventricles. The connective tissue is of the loose type originating in the pia mater. No qualitative morphological differences were observed between the choroid plexuses of the analyzed groups **(Fig.1).**

**Fig. 1.**
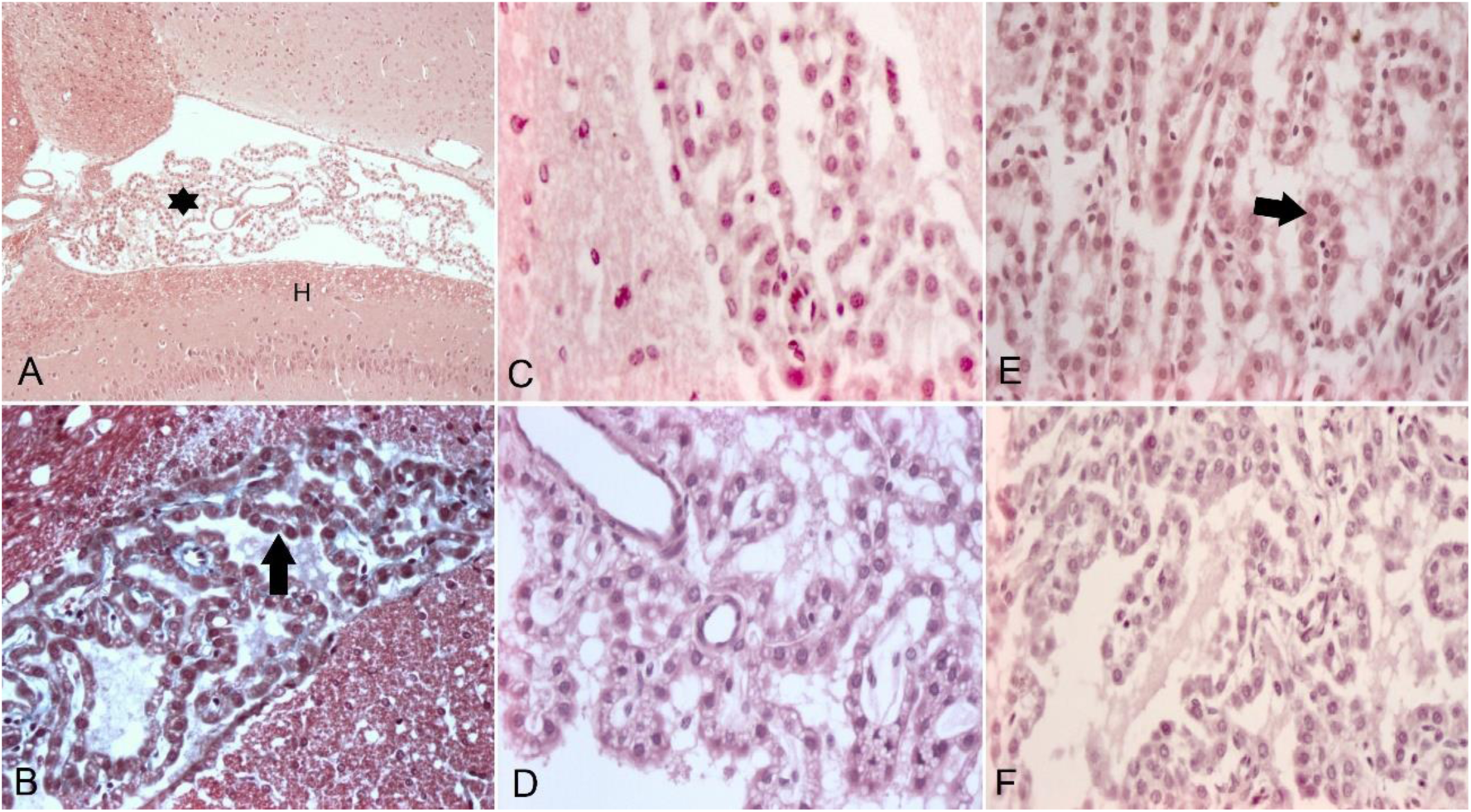
Photomicrographs of the rat brain. **A. Control group**. Observe cerebral ventricle with choroid plexus (*) and hippocampus (H). H.E 5X. **B**. **Control group**. Evidence of the choroid plexus composed of tufts of simple cubic epithelium (arrow). Masson trichrome 40X. **C**. **UChA Group** H. E. 40 X**. D. Control SM Group** H. E. 40 X. **E. UChB Group.** Observe choroid plexuses with simple cubic epithelium. **F. UChB SM group** H. E. 40.

### Morphometry

Table 1 shows a tendency towards epithelial atrophy of the choroid plexuses of UChA and UChB animals when compared to the control group. Epithelial atrophy was observed with a significant difference in the UChB MS group when compared to the Wistar group.

**Table 1.** Mean and standard deviation of epithelial height according to group and stress.

| Group | Stress |  |
| --- | --- | --- |
|  | Absent | Present |
| Control | 8,663 ± 0,515 aA* | 8,689 ± 0,501 aB |
| UChA | 8,449 ± 0,308 aA | 8,408 ± 0,328 aAB |
| UChB | 8,434 ± 0,152 aA | 8,174 ± 0,331 aA |

\* Lower letters: comparison of the fixed experimental condition and group
Capital letters: comparison of groups fixed to the experimental condition

The area of the choroid plexus epithelium of UChB animals was significantly larger when compared to the UChB MS. UChB MS showeds the smallest epithelial area **(Table 2).**

**Table 2.** Mean and standard deviation of the epithelium area according to group and stress.

| Group | Stress |  |
| --- | --- | --- |
|  | Absent | Present |
| Control | 25,313 ± 4,843 aA* | 25,583 ± 3,808 aA |
| UChA | 24,423 ± 2,828 aA | 25,654 ± 2,455 aA |
| UChB | 27,173 ± 1,501 bA | 23,714 ± 2,173 aA |

It also shows statistically significant results between the UChB and UChB MS groups for the epithelial perimeter of the choroid plexuses. UChB MS had smaller perimeters when compared to UChB animals **(Table 3).**

**Table 3.** Mean and standard deviation of epithelial perimeter according to group and stress.

| Group | Stress |  |
| --- | --- | --- |
|  | Absent | Present |
| Control | 18,899 ± 1,774 aA* | 19,014 ± 1,357 aA |
| UChA | 18,354 ± 1,309 aA | 18,930 ± 0,862 aA |
| UChB | 19,520 ± 0,551 bA | 18,178 ± 0,811 aA |

### Immunohistochemistry

Reactions of brain slides from animals from all groups showed strongly positive reactions for IGF-I in the choroid plexus. Strong positive reactions for this growth factor were also observed in similar blood vessels and pia maters in all groups **(Figs. 9-14).**

**Fig.**
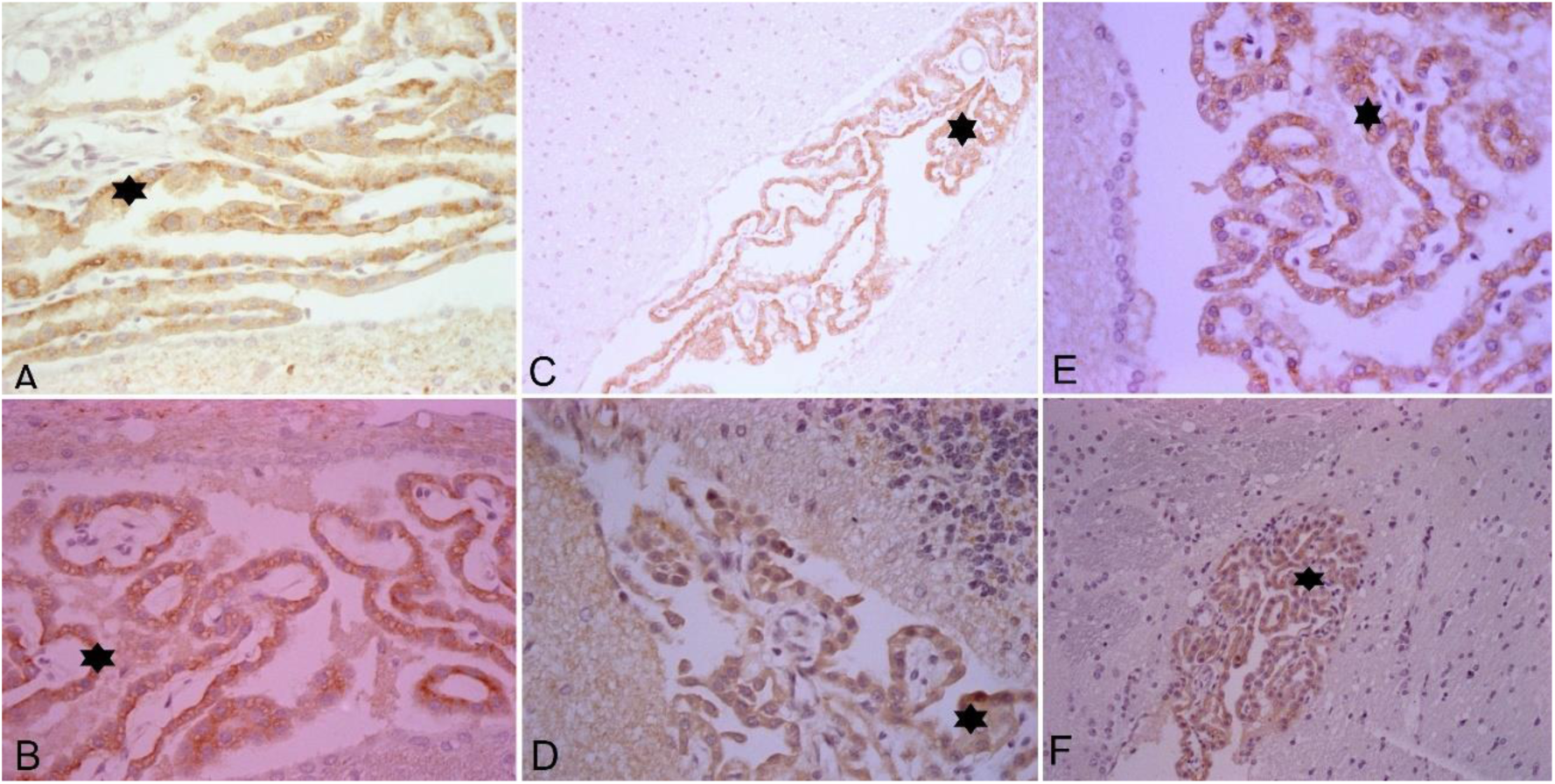
Photomicrographs of rat brain showing immunohistochemical reactions for IGF-I. **A**. Control and **B.** Control MS groups: shows strong positive reaction in the choroid plexuses (arrow) 40X. **C**. UChA and **D.** UChA MS Groups: observe a strong positive reaction for IGF-I in the choroid plexus (*) 20X and 40X. **E.** UChB and **F**. UChB MS Groups: Shows choroid plexus with strong reaction (*) 40X and 10X.

### Body Weight, Water, Ethanol and Feed consumption

#### Body weight

The experimental groups demonstrated the same pattern in relation to body mass, that is, greater gains with age. A higher mean was observed in SM control animals and a lower mean in UChB animals. Statistically significant differences were observed between animals in the control group and UChB SM at 120 days **(Table 4).**

**Table 4.** Mean and standard deviation of body weight (g) according to group and assessment moments (52th day and 120th day).

| Groups | Time of Assessment (age in days) |  | Statistical Test |
| --- | --- | --- | --- |
|  | 52 | 120 |  |
| UChA SM | 227,50±13,99dA | 364,00±15,05bcD | P<0,05 |
| UChA | 205,00±12,40bcA | 371,07±18,41bcE | P<0,05 |
| UChB SM | 193,57±28,98abA | 350,00±38,23 bD | P<0,05 |
| UChB | 183,57±29,18aA | 315,71±49,61aB | P<0,05 |
| Control SM | 219,64±20,33cdA | 398,21±37,29cD | P<0,05 |
| Control | 229,64±10,28dA | 385,00±33,51bcC | P<0,05 |
| Statistical Test Result | P<0,01 | P<0,001 | P<0,001 |

#### Water

**Table** 5 shows that water consumption in the UChA and UChB groups was lower when compared to the animals in the control group. Maternal separation decreased consumption in the UChB group and increased consumption in the UChA and control groups.

**Table 5.**
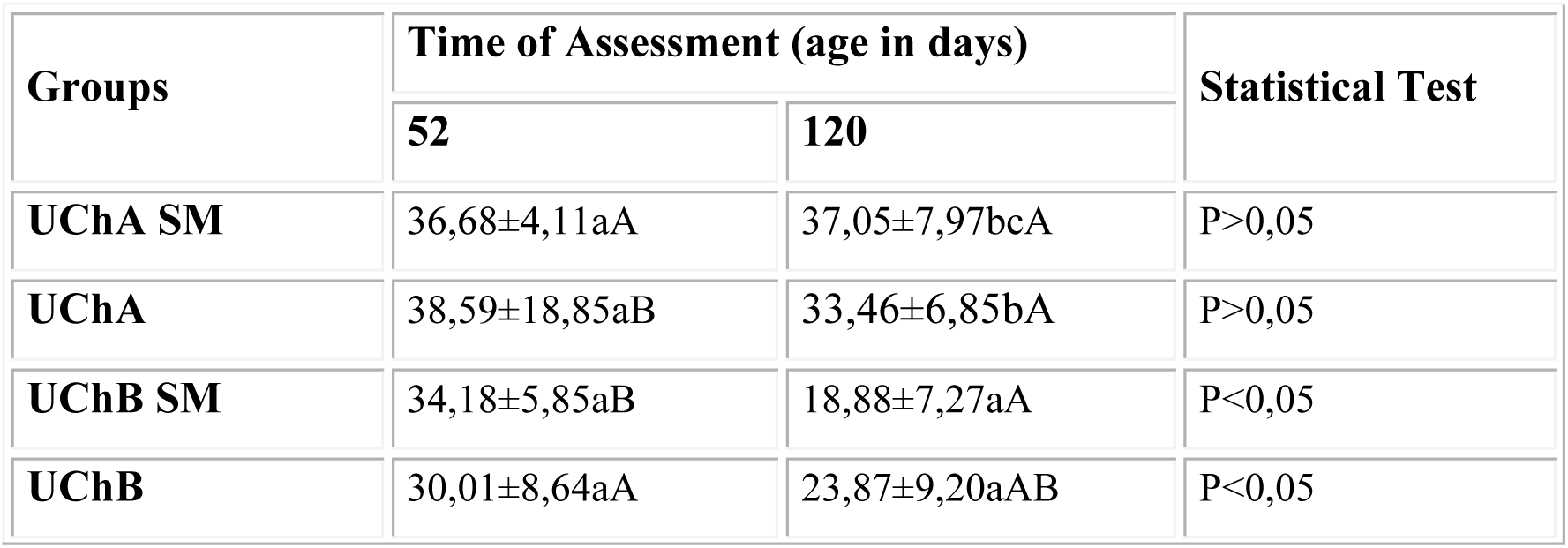
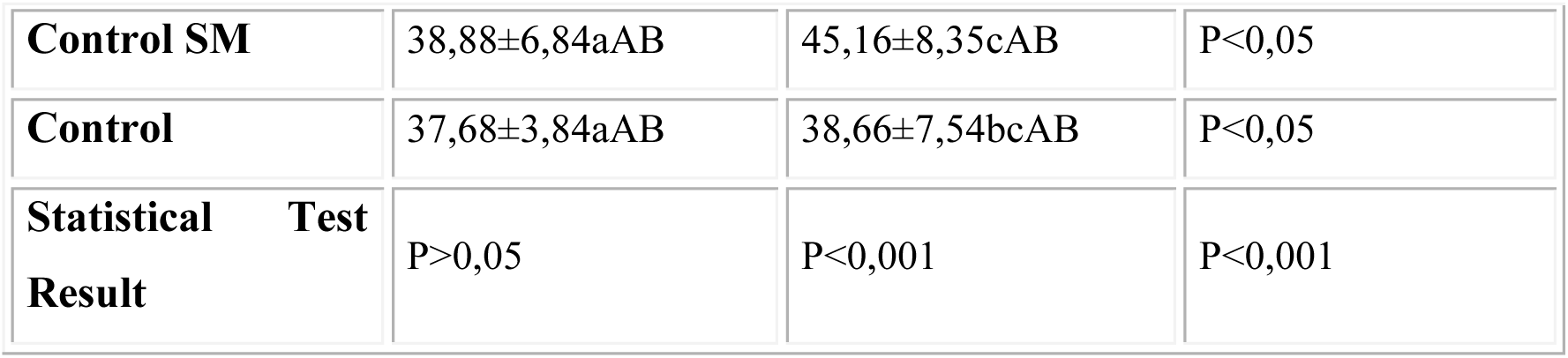
Mean and standard deviation of water consumption (ml) according to group and moment of assessment.

### Ethanol and Feed consumption

UChB animals showed statistically significant higher ethanol intake when compared to animals in the UChA group. Ethanol consumption showed different behavior between the UChA and UChB varieties in the face of maternal separation. The UChA group decreased consumption while the UChB group increased ethanol consumption **(Table 6).** The average feed consumption of animals in the UChB group was lower when compared to the Control and UChA groups. Feed consumption did not change with age and in the face of maternal separation. **(Table 7).**

**Table 6.** Mean and standard deviation of alcohol consumption (ml) according to group and moment of assessment.

| **Groups** | **Time of Assessment (age in days)** |  | **Statistical Test** |
| --- | --- | --- | --- |
|  | **73** | **120** |  |
| **UChA SM** | 4,71±1,71aA | 8,21±3,60aB | P<0,05 |
| **UChA** | 4,67±1,01aA | 11,52±6,35aB | P>0,05 |
| **UChB SM** | 15,12±4,20cA | 27,40±9,18cC | P<0,05 |
| **UChB** | 14,85±4,90cA | 21,64±9,60bcA | P>0,05 |
| **Statistical Test Result** | P<0,001 | P<0,001 | P<0,001 |

**Table 7.** Mean and standard deviation of feed consumption (g) according to group and moment of evaluation.

| **Groups** | **Time of Assessment (age in days)** |  | **Statistical Test** |
| --- | --- | --- | --- |
|  | **52** | **120** |  |
| **UChA SM** | 24,59±2,47cA | 28,11±3,48bAB | P<0,05 |
| **UChA** | 23,33±3,58bcA | 29,32±2,71bA | P>0,05 |
| **UChB SM** | 20,36±6,34abA | 22,51±5,26aA | P>0,05 |
| **UChB** | 18,57±3,19aA | 21,95±5,72aA | P>0,05 |
| **Control SM** | 22,98±3,76abcA | 25,95±2,91abA | P>0,05 |
| **Control** | 23,21±4,55bcA | 26,66±5,58abA | P>0,05 |
| **Statistical Test Result** | P>0,05 | P<0,001 | P<0,001 |

### Hormone Analysis

The plasma corticosterone of animals in the control group SM was significantly lower when compared to the control group. The UChA and UChB animals had lower corticosterone compared to the Wistar group. Maternal separation caused a statistically significant increase in corticosterone in these alcoholic animals. The FSH of the Wistar and UChB groups was lower when compared to neonatal maternal separation. The UChA group showed no difference. The LH of the Wistar group in the face of maternal separation was significantly lower. On the other hand, maternal separation increased the LH of UChA and UChB alcoholic animals. The testosterone hormone of the UCh groups were significantly lower when compared to the Wistar control. Maternal separation did not cause changes in the UCh groups, however, it reduced testosterone in the Wistar group. There was no change in estrogen between the groups, nor in relation to the experimental conditions **(Table 8).**

**Table 8.** Mean and standard deviation of hormones according to group and experimental condition.

| **Groups** | **Hormones** |  |  |  |  |
| --- | --- | --- | --- | --- | --- |
|  | **FSH** | **LH** | **Testosterone** | **Estrogen** | **Corticosterone** |
| **Control** | 9,9±3,6aA | 0,7±0,2bB | 1,0±0,4bB | 13,8±2,3aA | 57,4±16,4bB |
| **Control SM** | 6,9±2,7aA | 0,2±0,1aA | 0,7±0,3aB | 12,4±1,2aA | 18,6±11,7aA |
| **UChA** | 11,6±2,9Aa | 1,0±0,5aB | 0,3±0,1aA | 13,6±4,1aA | 10,0±4,4aA |
| **UChA SM** | 11,4±2,8aB | 2,2±1,1bB | 0,5±0,2aAB | 12,0±0,4aA | 28,6±13,5bA |
| **UChB** | 10,9±2,3aA | 0,1±0,1aA | 0,4±0,1aA | 12,2±2,9aA | 37,3±11,1aB |
| UChB SM | 8,5±2,8aAB | 0,7±0,3bA | 0,3±0,1 aA | 11,1±1,6aA | 67,2±24,8bB |

## DISCUSSION

The gain in body mass showed the same pattern between the experimental groups, with greater gains with age. Ethanol ingested in greater quantities in the UChB group decreases body mass gain with age. Maternal separation (MS) enabled increased mass gain in this group. Bo *et al*. (1982) described greater increase in body weight in rats in the control and isocaloric groups than in animals treated with ethanol. Similar results were described by Lee & Rivier (1993) and Cagnon *et al*. (1996). Contrasting data were published by Willis *et al*. (1983) in mice and Martinez *et al*. (2001) in Wistar rats who did not observe significant differences between the average body weights of the groups studied. Our experiment showed lower feed consumption in the UChB group and higher in the UChA group. Maternal separation did not change the consumption of this variable. Several studies show a decrease in body weight in animals treated with ethanol (Cagnon *et al*., 1996; Oliveira & Ferreira, 1987). Ratcliffe (1972) commented that the decrease in weight gain may be caused by irritation caused in the gastrointestinal tract, reduction in the production of endogenous testosterone, with excessive conversion of testosterone into estradiol and increased lipid oxidation. In the present study, low testosterone levels coincide with body mass data, with Wistar>UChA≥UChB.

Ethanol consumption showed different behavior between the UChs varieties about maternal separation. The UChA decreased while the UChB group increased ethanol consumption at the end of the experiment. Previous studies revealed that stressful events during rodents’ childhood, such as maternal separation, favor ethanol consumption, and dependence (Gilmer & Mckinney, 2003; Filarowska-Jurko *et al*. 2022). Ethanol intake in UCh lineages is mainly related to the difference in the ability to metabolize acetaldehyde by the ALDH2 enzyme (Quintanilla & Tampier, 2003). It is suggested that during the maternal separation procedure there was no sensitization of the acetaldehyde metabolization mechanisms, preserving the characteristic consumption of the UChA lineage. However, possible compensatory mechanisms should not be excluded, such as metabolism, genetic vulnerability, lifestyle, sex, nutritional factors, time, and amount of ethanol use. Another relevant factor is related to differences in behavior of UCh females during the Maternal Separation period. UChA females showed intense maternal care behavior, called alpha behavior. UChB females showed maternal behavior of low care for their offspring, called beta. Puppies from mothers with a high level of care have less fearful behavior when faced with new situations in adulthood, with a mild response of the hypothalamic-pituitary-adrenal axis to stress, when compared to puppies from mothers with a low level of maternal care (Francis *et al*. 1999). In other words, maternal behavior can modulate the development of individual HPA axis responses and, thus, influencing behavioral responses to stressors in adults (Caldji *et al*., 1998).

Maternal separation showed a positive correlation with the release of serum corticosterone in adulthood, corroborating the literature (Biagini *et al*., 1998; Knuth *et al*. 2005). This occurs due to the decreased efficiency of corticosterone feedback regulation in the hypothalamus, which makes the HPA axis hyper-responsive to stress (Biagini *et al*., 1998). It was verified in the experiment that ethanol ingested in adulthood is an aggressive agent that upregulated the activity of the HPA axis in the release of corticosterone in rats subjected to maternal separation. Rats subjected to maternal separation that did not ingest ethanol showed a decrease in serum corticosterone values. The inability to metabolize acetaldehyde due to chronic ethanol intake, as well as its accumulation, leads to organic toxicity, which stimulates the activity of the HPA axis (Haddad, 2004).

Plasma testosterone levels showed no difference between strains and groups, but when compared with the literature, it remained below normal standards. Low plasma testosterone levels were observed in ethanol-treated rats (Oliva *et al*. 2006). Anderson Jr *et al*. (1987) reported that chronic ethanol exposure exerts direct effects on all levels of the HHG axis, reducing luteinizing hormone and testosterone levels. Kim *et al*. (2003) indicated that ethanol acts by inhibiting the release of GnRH, resulting in a decrease in FSH, LH and consequently testosterone. However, the present study showed higher plasma LH content in the SM group of the UChA lineage compared to the control and the other SM groups. The inhibition of LH synthesis may be due to the direct effect of ethanol on the gonads. In short, data in the literature seem to reveal that there are ideal levels of serum corticosterone for the adequate production and release of testosterone. Although serum testosterone levels were not altered in our experiment, changes in tissue testosterone and paracrine relationships should not be ignored, since corticosterone receptors were found to be present in the cells of the seminiferous tubules and interstitium (Biagini & Pich, 2002).

The results of the present research showed a tendency towards epithelial atrophy in UCh animals and an increase in this atrophy due to maternal separation. Clair (1991) reported the presence of dilated brain ventricles in chronic alcoholics. Tirapelli *et al*. (2000) described ultrastructural changes in the choroid plexus of adult Wistar rats treated with sugar cane spirit. Thiamine deficiency and ethanol oxidation cause damage to the epithelial cells of the choroid plexus. These cells are responsible for maintaining glutamate homeostasis. With its reduced functioning, glutamate is not metabolized. It is known that the choroid plexuses are fundamental structures in the creation and maintenance of homeostasis in the central nervous system. Possible effects on the morpho functional integrity in the choroid plexuses can alter the blood-CSF barrier and indirectly the blood-brain barrier, inducing serious disorders in the CNS (Nixon 2008).

The results showed strong expression of IGF-I in blood vessels, choroid plexus and pia mater in the animals analyzed. The literature shows conflicting results regarding this growth factor, stimulating new methodological approaches. Sonntag *et al*. (1999), the expression of IGF-I in the brain of rats was significant in arteries, arterioles, and meninges. The gene expressions of this growth factor did not change with age, while the protein expression decreased, suggesting deficiencies in transport across the blood-brain barrier. Chung *et al*. (2002) investigated the relationship between age and IGF-I receptors in the cerebral cortex and hippocampus, showing a significant increase in immunoreactivity of IGF-I receptors. On the other hand, IGF-I immunoreactivity was absent in the hippocampus and dentate gyrus of the Mongolian gerbil (Hwang *et al*. 2004). In groups subjected to the stress of maternal separation, there was a decrease in the expression of transiterin, as well as IGF-II. This indicates that the choroid plexus has an important role in susceptibility to stress, in addition to neuronal and glial roles. (Kohda *et al*. 2006). De La Monte *et al*. (2008) cited a reduction in the expression of insulin, IGF-I and IGF-II and their receptors in the cerebellum of human alcoholics. Chronic prenatal ethanol exposure decreased mRNA expression of IGF-1, IGF-1R, and IGF-2 (Dobson *et al*. 2014). There was greater expression of IGFR-1 in the UChB groups with strong labeling in Purkinje cells (Martinez *et al*. 2018). The toxic effects of ethanol are not uniform: some brain regions and cell types are more affected than others. Thus, it is understood that ethanol can interfere with all stages of brain development with cognitive, emotional, and behavioral impairments (Guerri, 2002).

## CONCLUSIONS

Maternal separation (MS) reversed the process triggered by alcoholism in the UChB body mass group. There was a positive association in the UChB group and a negative association in the UChA between water consumption and MS. There was a negative association between ethanol consumption and MS. MS caused some hormonal imbalance in UCh animals. Maternal separation potentiated epithelial atrophy of the choroid plexus in UChA and UChB animals.

